# Immunometabolic remodeling of perivascular adipose tissue in murine lupus: implications for lupus vasculopathy

**DOI:** 10.64898/2026.05.18.726104

**Authors:** Lingling Liu, David Kim, Yong Zhang, Brandee Goo, Xin Xiong, Ting Zhang, Qimei Han, Shitong Wu, Qingkang Lv, Mourad Ogbi, Tianxiang Hu, Hanping Wu, Vijay S. Patel, Dominic Gallo, Rachard Lee, Ha Won Kim, David Stepp, David Fulton, J Michelle Kahlenberg, Laura Carbone, Klaus Ley, Brian H Annex, Neal L Weintraub, Hong Shi

## Abstract

**Background:** Patients with systemic lupus erythematosus (SLE) face markedly increased cardiovascular disease (CVD) risk driven by mechanisms beyond traditional risk factors. Thoracic aortic perivascular adipose tissue (tPVAT) is dysfunctional in lupus and exacerbates endothelial dysfunction, yet the molecular basis of this dysfunction remains poorly defined.

**Methods:** Integrated multi-omics profiling, including bulk RNA-seq, untargeted proteomics, lipidomics, and high-dimensional spectral flow cytometry, was performed on tPVAT from 15-week-old MRL/lpr mice (active lupus, n = 4–6) and MRL control mice (n = 5–6). Adipogenic differentiation capacity of tPVAT adipose stromal and progenitor cells (ASPCs) from MRL/lpr was assessed by Oil Red O staining at 5 (pre-dieasea) and 15 weeks (active disease), with subcutaneous ASPCs used as depot controls.

**Results:** Transcriptomic profiling of tPVAT from MRL/lpr mice identified 2,742 upregulated and 1,494 downregulated genes (adjusted p < 0.001, |log2FC| > 1), with strong activation of interferon, IL6-JAK-STAT3, and TNFA signaling pathways together with suppression of fatty acid metabolism, oxidative phosphorylation, and adipogenic pathways. Proteomic and lipidomic analyses were concordant, revealing broad downregulation of mitochondrial bioenergetic machinery, depletion of cardiolipin and acylcarnitines, and enrichment of ceramide phosphoinositols and lysophosphatidylcholines. Cardiolipin strongly correlated with the mitochondrial/metabolic protein module (r = 0.95) and inversely with the immune/inflammatory protein module (r = −0.92). Spectral flow cytometry confirmed marked CD45+ leukocyte infiltration dominated by T cells, together with a significantly reduced Treg/CD4+ ratio indicating loss of local immunoregulatory balance. ASPCs derived from PVAT of 15-week-old MRL/lpr mice exhibited impaired white and beige adipogenic differentiation, while APCs from PVAT of 5-week-old MRL/lpr mice, and from subcutaneous adipose tissues of 15-week-old MRL/lpr mice, had normal white and beige differentiation, consistent with an acquired, depot-specific, disease-stage-dependent progenitor defect in PVAT of MRL/lpr mice.

**Conclusions:** Lupus tPVAT undergoes a concordant cross-platform molecular reprogramming of mitochondrial bioenergetic genes coupled with establishment of an interferon-dominant immune niche and acquired loss of ASPC adipogenic capacity. These findings provide a molecular framework for lupus PVAT dysfunction and identify restoration of mitochondrial function, suppression of interferon-driven inflammation, and renewal of progenitor differentiation as potential therapeutic strategies for lupus vasculopathy.

## Introduction

Systemic lupus erythematosus (SLE) is a chronic autoimmune disorder characterized by profound immune dysregulation and multiorgan involvement, disproportionately affecting women of childbearing age [1]. Despite improvements in disease management, cardiovascular disease (CVD) remains a leading cause of mortality in SLE, with young women carrying a CVD risk reported to be up to 50-fold higher than age-matched controls [2–9]. This excess risk cannot be fully explained by traditional Framingham risk factors, implicating disease-specific mechanisms, such as accrued organ damage, chronic systemic inflammation, and Type I/II interferon-driven immune activation, in the pathogenesis of premature atherosclerosis in lupus [9–11].

Perivascular adipose tissue (PVAT) is a metabolically active depot surrounding most large blood vessels that plays an important role in regulating vascular homeostasis. Under healthy conditions, thoracic aortic PVAT (tPVAT) resembles thermogenic brown or beige adipose tissue and exerts vasoprotective effects through adipokine secretion (e.g., adiponectin), promotion of fatty acid oxidation, and Ucp1-mediated thermogenesis [12]. In the setting of cardiovascular risk factors, tPVAT undergoes phenotypic switching, losing thermogenic capacity, acquiring a white-like morphology, and becoming infiltrated by immune cells, thereby promoting endothelial dysfunction, oxidative stress, and vascular inflammation through an “outside-in” mechanism [13–15]. Recent single-cell and spatial omics studies have further highlighted PVAT as an immunologically active organ, with depot-specific stromal and immune cell niches that shape vascular tone and inflammatory responses [16–18].

Clinical evidence increasingly identifies PVAT as a central player in lupus-related CVD. Women with SLE exhibit greater volumes and higher densities of tPVAT (which correlates with heightened inflammation) compared to healthy controls, and PVAT density independently associates with aortic calcification, a marker of subclinical atherosclerosis, after adjusting for age, BMI, and circulating inflammatory markers [19, 20]. PVAT characteristics have also been linked to adverse metabolic parameters, suggesting that PVAT is a core component of the lupus cardiometabolic phenotype [21]. Experimentally, tPVAT from lupus-prone MRL/lpr mice is dysfunctional, exhibiting adipocyte hypertrophy, whitening, collagen deposition, and immune cell infiltration, together with reduced expression of adipogenic (Pparg) and thermogenic (Ucp1) genes and elevated pro-inflammatory cytokines and chemokines [22]. Critically, this dysfunctional PVAT exacerbates endothelial dysfunction in mice with lupus, supporting a direct pathogenic role for inflamed PVAT in lupus-associated CVD [22]. In parallel, serum metabolomic and lipidomic studies in SLE have identified phospholipid and sphingolipid signatures—including lysophosphatidylcholines and ceramides—that associate with systemic inflammation and subclinical atherosclerosis, suggesting a role for immunometabolic lipid remodeling in lupus CVD [23].

However, the molecular and cellular mechanisms driving PVAT dysfunction in lupus remain poorly defined. In particular, the transcriptomic, proteomic, and lipidomic landscape of PVAT in lupus, and how these converge to impair the tissue’s vascular-protective functions, has not been systematically investigated. Furthermore, it is unknown whether the PVAT inflammatory niche in lupus may compromise the regenerative and thermogenic capacity of resident adipose stem progenitor cells (ASPCs), which are essential for adipocyte renewal and maintenance of beige adipocyte identity.

In the present study, we utilized an integrated multi-omics approach, combining bulk RNA-seq, untargeted proteomics, lipidomics, and high-dimensional spectral flow cytometry, to characterize the tPVAT from 15-week MRL/lpr (lupus prone) and MRL control mice. We demonstrate sweeping, cross-platform remodeling defined by a concordant diminution of mitochondrial bioenergetic programs, including depletion of cardiolipin and loss of fatty acid oxidative capacity, together with remodeling of the lipidome toward pro-inflammatory, lipotoxic species. These metabolic changes occur in parallel with the establishment of a chronic, T cell–dominated, and interferon-rich immune niche within the perivascular compartment. Importantly, we provide functional evidence that this environment imposes an acquired, depot-specific defect in ASPC adipogenic differentiation. Together, these findings provide a mechanistic, immunometabolic framework for how PVAT loses its vascular-protective identity and contributes to accelerated vasculopathy in lupus.

## Materials and Methods

### Animals

Female MRL/MpJ (MRL, stock #000486) and MRL/MpJ-Faslpr/J (MRL/lpr, stock #000485) mice were purchased from The Jackson Laboratory (Bar Harbor, ME) and maintained under specific pathogen-free conditions with a 12-hour light/dark cycle and ad libitum access to food and water. Only female mice were used in the experiments described here. MRL and MRL/lpr mice were harvested at 15 weeks of age, corresponding to active lupus disease in MRL/lpr mice, as confirmed by proteinuria and elevated anti-dsDNA antibodies (22). All animal care and experimental protocols complied with the National Institutes of Health Guide for the Care and Use of Laboratory Animals and relevant ethical regulations, and were approved by the institutional Animal Care & Use Committee of Augusta University (AU).

### Tissue harvesting

At age of 15 weeks, tPVAT was carefully dissected from the thoracic aorta following perfusion with phosphate-buffered saline. Tissue was weighed and divided for downstream applications including bulk RNA sequencing, mass spectrometry-based proteomics and lipidomics, flow cytometry, and adipose stromal/progenitor cell (ASPC) isolation.

### Bulk RNA Sequencing and Differential Expression Analysis

Total RNA was extracted from tPVAT using the RNeasy Lipid Tissue Mini Kit (Qiagen) according to the manufacturer’s instructions. RNA integrity and purity were assessed using a Bioanalyzer (Agilent). Library preparation and sequencing were performed by GENEWIZ (Azenta Life Sciences, project 30-548589527) on an Illumina platform with 150-bp paired-end reads. Nine tPVAT samples (MRL n=5, MRL/lpr n=4; one MRL/Lpr sample excluded due to QC failure) yielded a mean of 61.4 million reads per sample (range 51.2–67.7 million; mean quality score ≥37.94; >90.8% bases at Q≥30). Due to quality degradation in the 100–150 bp region, reads were trimmed to 100 bp prior to alignment. Trimmed reads were pseudoaligned to the mouse reference transcriptome (GRCm39, Ensembl 104) using Kallisto (v0.46.1) with 100 bootstrap samples, and transcript-level abundance estimates were imported and summarized to gene level using tximport. Differential expression analysis was performed with DESeq2 (v1.48.1) in R, and genes with an adjusted p-value <0.001 and |log₂ fold change|>1were considered differentially expressed. Gene set enrichment analysis (GSEA) was carried out using the fgsea package with Hallmark gene sets from MSigDB.

### Proteomics and Lipidomics

Untargeted proteomics and lipidomics were performed by Dalton Bioanalytics (Los Angeles, CA) using the One Shot Multi Omics platform on tPVAT from MRL (n = 5) and MRL/lpr (n = 5) mice. This platform enables simultaneous extraction of proteins and lipids from a single sample aliquot, ensuring high biological consistency across platforms. For proteomics, proteins were digested with trypsin and analyzed by liquid chromatography-tandem mass spectrometry (LC-MS/MS). For lipidomics, lipids were extracted and analyzed by LC-MS using both positive and negative electrospray ionization modes to maximize lipidome coverage. Differential abundance was determined using the vendor-provided lm3.thor_lupus linear modeling framework, which applies a contrast-based model (MRL/lpr vs MRL) to each molecular feature. Proteins and lipids with an adjusted p value < 0.05 and |log2 fold change| > 0.5 were considered statistically significant. Multi-omics integration, including pathway concordance analysis and module scoring, was performed in R. To summarize coordinated changes across multiple proteins, mitochondrial/metabolic and immune/inflammatory protein modules were constructed by calculating the mean log-transformed protein abundance across selected proteins within each category. The mitochondrial/metabolic protein module comprised concordantly downregulated proteins identified in the integrated analysis (*Ckmt2, Cox17, Slc25a1, Sfxn1, Pdha1, Ldha, Cox6b1, Pkm,* and *Echdc1*). The immune/inflammatory protein module comprised concordantly upregulated immune-associated proteins (*Igkc, H2-Aa, Stat1, Samhd1, Cybb, Coro1a, Rac2, Ptpn6, Pld4,* and *Arhgdib*). Pearson correlations were calculated between lipid class summary scores and protein module scores across all samples.

### Spectral Flow Cytometry

tPVAT was enzymatically dissociated into single-cell suspensions using an Adipose Tissue Dissociation Kit (Miltenyi Biotec) according to the manufacturer’s protocol. Cells were incubated with Fc block (anti-CD16/32) to prevent nonspecific binding, followed by surface staining with the Cytek 24-Color Mouse Immunoprofiling Panel (Cytek Biosciences) for 30 minutes at 4°C. Viability was assessed using a fixable viability dye (Cytek Biosciences). Data were acquired on a five-laser Cytek Aurora spectral flow cytometer. Raw spectral data were unmixed using SpectroFlo software (Cytek Biosciences) with single-stain reference controls and unstained cells for autofluorescence subtraction. Final population gating was performed in SpectroFlo, and downstream analysis was conducted in R (version 4.5). Cell counts were normalized to the initial tissue weight and expressed as cells per gram. Statistical comparisons for cell-count data were performed using unpaired two-tailed Student’s t-tests on log10-transformed values to improve normality, and p-values <0.05 were considered statistically significant.

### ASPC Isolation and In Vitro Adipogenesis

ASPCs were isolated by negative magnetic selection using the Miltenyi MACS system to deplete CD45⁺, CD31⁺, and Ter119⁺ cells, yielding a CD45⁻CD31⁻Ter119⁻ stromal fraction. For each independent biological replicate (N=3), tPVAT from three mice was pooled to ensure sufficient and representative progenitor populations (total n=9 mice per genotype), and differentiation assays were performed in technical triplicate. ASPCs were induced toward white or beige adipogenic differentiation using standard adipogenic induction cocktails. After 10 days of differentiation, lipid accumulation was quantified by Oil Red O (ORO) staining; ORO was extracted with isopropanol and absorbance measured at 490 nm, then normalized to total protein content determined by BCA assay. Representative phase-contrast microscopy images were acquired at 10× magnification (scale bar, 150 µm).

### Statistical Analysis

Data are presented as mean ± standard error of the mean (SEM). Comparisons between two groups were generally performed using unpaired two-tailed Student’s t-tests; for flow cytometry-derived cell counts, tests were applied to log10-transformed values to improve normality. For adipogenesis assays, unpaired two-tailed Student’s t-tests were performed separately for white and beige differentiation conditions, as indicated in the figure legends. A p value < 0.05 was considered statistically significant. All statistical analyses and data visualization were performed in R (version 4.5) using RStudio.

## Results

### Lupus drives broad transcriptomic reprogramming of tPVAT, marked by inflammatory activation and loss of thermogenic and stromal matrix programs

To define the molecular landscape of thoracic perivascular adipose tissue in lupus, we performed bulk RNA sequencing on tPVAT from 15-week-old MRL mice (n = 5) and MRL/lpr mice (n = 4). Principal component analysis demonstrated clear separation between groups along PC1, which accounted for 38.98% of the total variance, indicating a strong disease-associated transcriptional signature in tPVAT from MRL/lpr mice (Figure 1A)[22]. Differential expression analysis identified 2,742 upregulated and 1,494 downregulated genes in tPVAT from MRL/lpr mice relative to MRL mice (adjusted p<0.001, |log₂FC|>1), consistent with extensive transcriptional reprogramming of the depot (Figure 1B). Hallmark pathway enrichment further revealed an enrichment of inflammatory pathways including Allograft Rejection, Interferon Gamma Response, Interferon Alpha Response, IL6-JAK-STAT3 Signaling, Inflammatory Response, and TNFA Signaling via NF-κB, together with suppression of core metabolic pathways such as Fatty Acid Metabolism, Oxidative Phosphorylation, Adipogenesis, Peroxisome, Cholesterol Homeostasis, and Bile Acid Metabolism (Figure 1C) [24]. Consistent with this loss of metabolic fitness, several thermogenic and mitochondrial genes were reduced in tPVAT from MRL/lpr mice, including *Ucp1, Cidea, Cox8b, Elovl3, Elovl5, Ndufa1, Opa1,* and *Mfn2* (Figure 1D)[25]. In contrast, Ucp2 was selectively upregulated, a pattern more consistent with a stress-adaptive, reactive oxygen species-buffering state than a canonical thermogenic program[26, 27]. Hierarchical clustering of representative inflammatory, interferon-responsive, and immune activation genes demonstrated consistent induction across biological replicates, including *Stat1, Irgm1, Irgm2, Il1b,* and *Tgtp1*, supporting an inflammatory remodeling of from MRL/lpr mice tPVAT (Figure 1E) [28–31].

**Figure 1.**
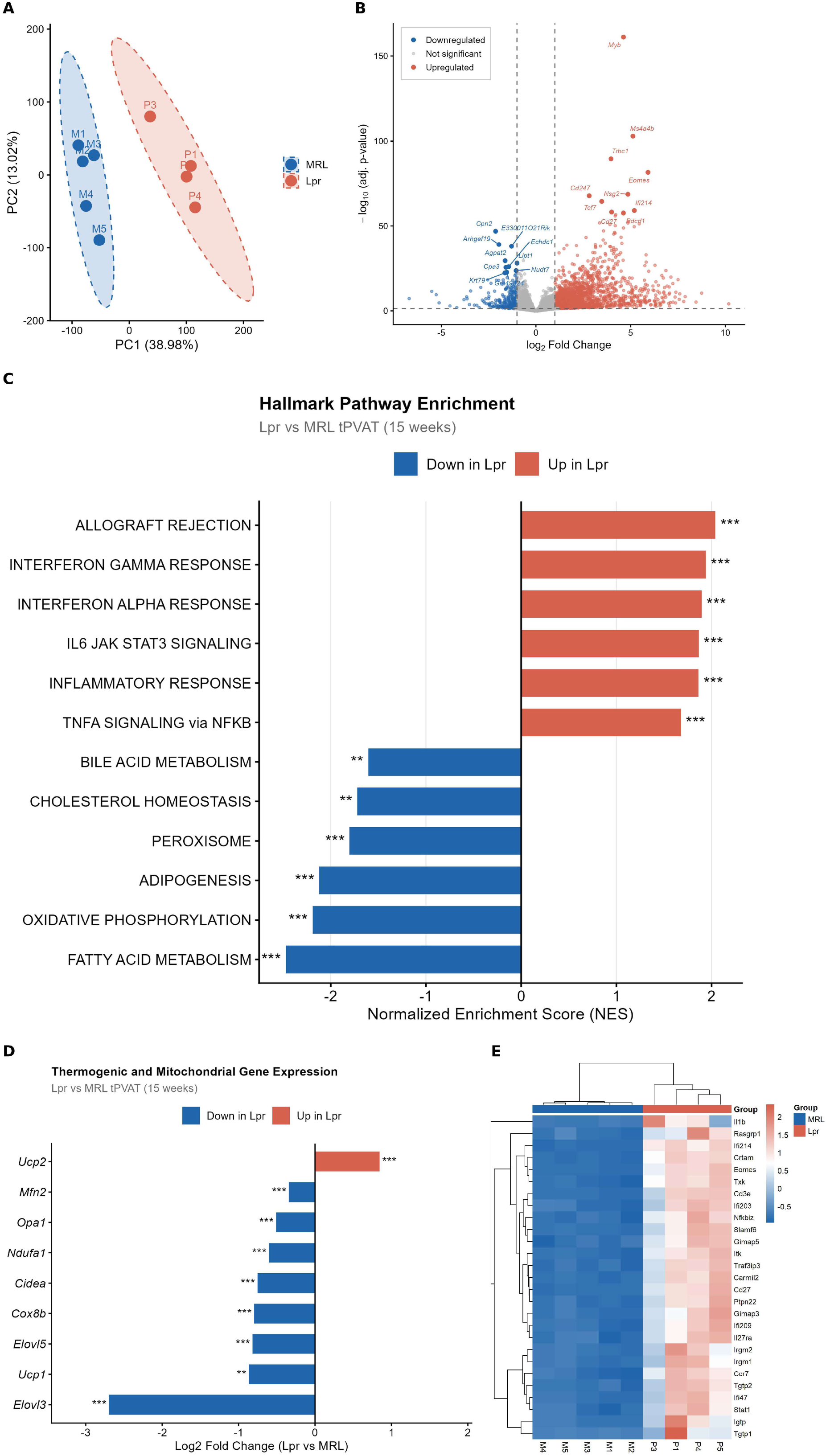
Lupus-driven metabolic collapse and inflammatory transformation of the thoracic perivascular adipose tissue (tPVAT) transcriptome. (A) Principal component analysis (PCA) of bulk RNA-seq data from tPVAT of 15-week-old MRL (control) and MRL/Lpr (lupus) mice (MRL, n = 5; MRL/Lpr, n = 4), demonstrating distinct transcriptomic clustering. (B) Volcano plot showing differentially expressed genes (DEGs) in MRL/Lpr versus MRL tPVAT (adjusted p < 0.001, |log2FC| > 1); red indicates upregulated genes and blue indicates downregulated genes. (C) Hallmark pathway enrichment analysis showing suppression of metabolic pathways, including Fatty Acid Metabolism, Oxidative Phosphorylation, and Adipogenesis, together with activation of inflammatory signaling pathways, including Interferon-γ Response, IL6-JAK-STAT3 Signaling, and TNFA Signaling via NF-κB. (D) Expression profile of selected thermogenic and mitochondrial genes, highlighting loss of brown/beige adipocyte identity with selective upregulation of the stress-responsive gene *Ucp2* in Lpr tPVAT. (E) Hierarchical clustering heatmap of representative inflammatory and interferon-responsive genes across individual biological replicates (z-score normalized expression).

To examine matrix-related changes, we generated a curated heatmap of fibrosis-associated structural genes, structural ECM genes, and inflammatory recruitment/remodeling genes (Figure S1). This analysis showed that tPVAT from MRL/lpr mice had lower expression of multiple stromal and matrix-organizing genes, including *Col6a1, Col6a2, Col1a1, Fbn1, Pcolce2, Lum, Col5a1, Col5a2, Col4a2, Dcn,* and *Pcolce* [32]. In parallel, inflammatory recruitment genes such as *Ccl8, Vcam1, Itgax,* and *Itgb2* were increased [33–35]. Notably, matrix metalloproteinase genes including *Mmp12, Mmp9, Mmp2,* and *Mmp19* were reduced rather than induced in tPVAT from MRL/lpr mice [36]. Together, these findings indicate that in MRL/lpr mice, tPVAT undergoes coordinated inflammatory reprogramming accompanied by loss of brown/beige-associated metabolic identity and disruption of homeostatic stromal matrix programs, rather than uniform activation of a classic fibrotic transcriptional response.

### tPVAT from MRL/lpr mice exhibits marked immune infiltration and impaired immunoregulatory balance

To define the cellular basis of this inflammatory transcriptomic shift observed in lupus tPVAT, we performed high-dimensional spectral flow cytometry on tPVAT from 15-week MRL (n=6) and MRL/lpr(n=6) mice. Total CD45⁺ leukocyte counts were significantly increased in tPVAT from MRL/lpr mice, indicating a substantial immune accumulation within perivascular niche (Figure 2A). Across major immune populations, T cells were the most prominently expanded compartment, while dendritic cells were also significantly increased. Macrophages were numerically higher in tPVAT of MRL/lpr mice, although this difference did not reach statistical significance (p = 0.054). B cells, NK cells, and monocytes likewise showed upward trends that were not statistically significant, whereas neutrophils were unchanged (Figure 2B). Within the T cell compartment, both CD4+ and CD8+ T cells were significantly elevated in PVAT from MRL/lpr mice, and Tregs were also increased in absolute number (Figure 2C). However, despite this increase in absolute Treg number, the Treg/CD4+ ratio was significantly reduced in tPVAT from MRL/lpr mice, indicating that regulatory T cell expansion did not keep pace with the broader accumulation of CD4+ T cells (Figure 2D) [37, 38]. Subset analysis further showed selective enrichment of CD44−CD62L− cells within both the CD4+ and CD8+ compartments, whereas naive, central memory, and effector memory subsets were not significantly altered (Figure 2E, F). Together, these findings indicate that in MRL/lpr mice, tPVAT undergoes pronounced immune remodeling characterized by T cell accumulation and relative loss of immunoregulatory balance.

**Figure 2.**
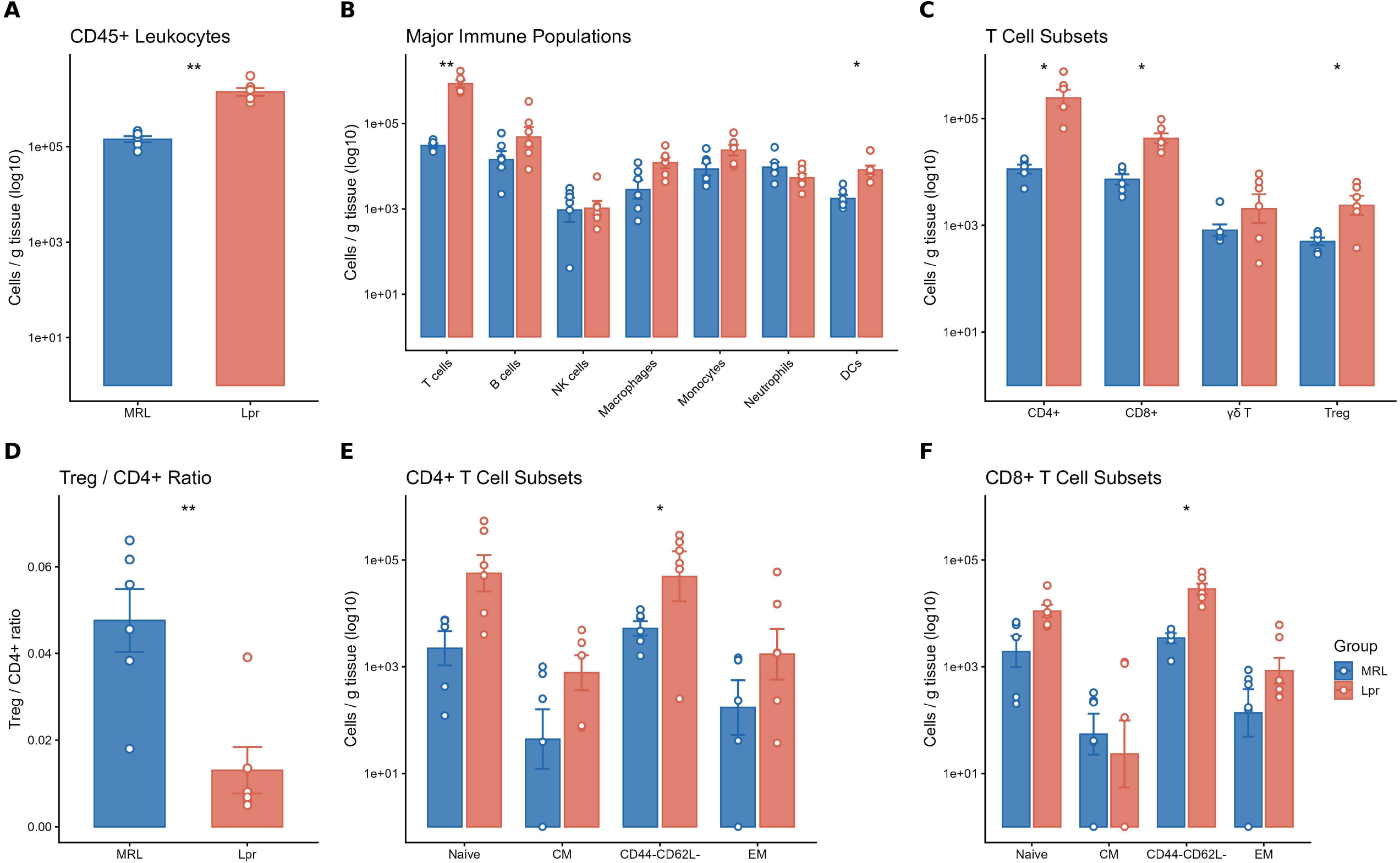
Spectral flow cytometry reveals marked immune infiltration and loss of immunoregulatory balance in lupus tPVAT. (A) Total CD45+ leukocyte counts per gram of tPVAT in 15-week-old MRL control (n = 6) and MRL/Lpr (n = 6) mice. (B) Major immune cell populations normalized to tissue weight, including T cells (CD3+TCRβ+), B cells (B220+CD19+), NK cells (CD49b+NK1.1+CD3−), macrophages (CD45+F4/80+CD11b+), monocytes, neutrophils (Ly6G+CD11b+), and dendritic cells (DCs; CD11c+MHC II+). (C) T cell subset counts per gram of tissue, including CD4+, CD8+, γδ T cells (CD3+TCRγδ+), and CD4+CD25^hi regulatory T cells (Tregs). (D) Ratio of Tregs (CD4+CD25^hi) to total CD4+ T cells, reflecting the relative abundance of regulatory cells within the CD4+ compartment. (E) CD4+ T cell subsets normalized to tissue weight: naïve (CD44−CD62L+), central memory (CM; CD44+CD62L+), CD44−CD62L− atypical/pre-effector-like cells, and effector memory (EM; CD44+CD62L−). (F) CD8+ T cell subsets normalized to tissue weight, using the same phenotypic definitions as in (E). Data are presented as mean ± SEM; each dot represents one biological replicate. Statistical comparisons were performed by unpaired t test on log-transformed values. *P < 0.05, **P < 0.01, ***P < 0.001, ****P < 0.0001.

### Proteomics and lipidomics reveal concordant mitochondrial dysfunction and lipidome remodeling in lupus tPVAT

To determine whether the transcriptional alterations in lupus tPVAT were reflected at the protein and lipid level, we performed untargeted proteomics and lipidomics on tPVAT from 15-week MRL (n=5) and Lpr (n=5) mice. Principal component analysis of both datasets showed clear group separation, indicating that lupus imposes distinct proteomic and lipidomic signatures on tPVAT (Figure 3A,B). Volcano plot analysis of the proteomic dataset identified increased abundance of immune-related proteins, including immunoglobulin-associated proteins and F13a1, together with reduced abundance of metabolic enzymes such as Fasn, Acly, and Acaca in Lpr tPVAT (Figure 3C). Proteomics-based pathway enrichment demonstrated broad suppression of mitochondrial and bioenergetic programs, including mitochondrial inner membrane organization, aerobic respiration, fatty acid β-oxidation, the TCA cycle, and mitochondrial translation, alongside enrichment of adaptive immune response and platelet-associated pathways (Figure 3D). Lipidomic analysis further revealed marked remodeling of the lipidome in PVAT from MRL/lpr mice, characterized by depletion of acylcarnitines, fatty acid esters of hydroxy fatty acids (FAHFAs), cardiolipin, hydroxy-PE species, and diacylglycerols, together with enrichment of ceramide phosphoinositols and lysophosphatidylcholines (LPCs) (Figure 3E). Together, these data indicate that tPVAT from MRL/lpr mice undergoes dysfunctional metabolic remodeling at the proteomic level, accompanied by accumulation of inflammatory and membrane-remodeling lipid species.

**Figure 3.**
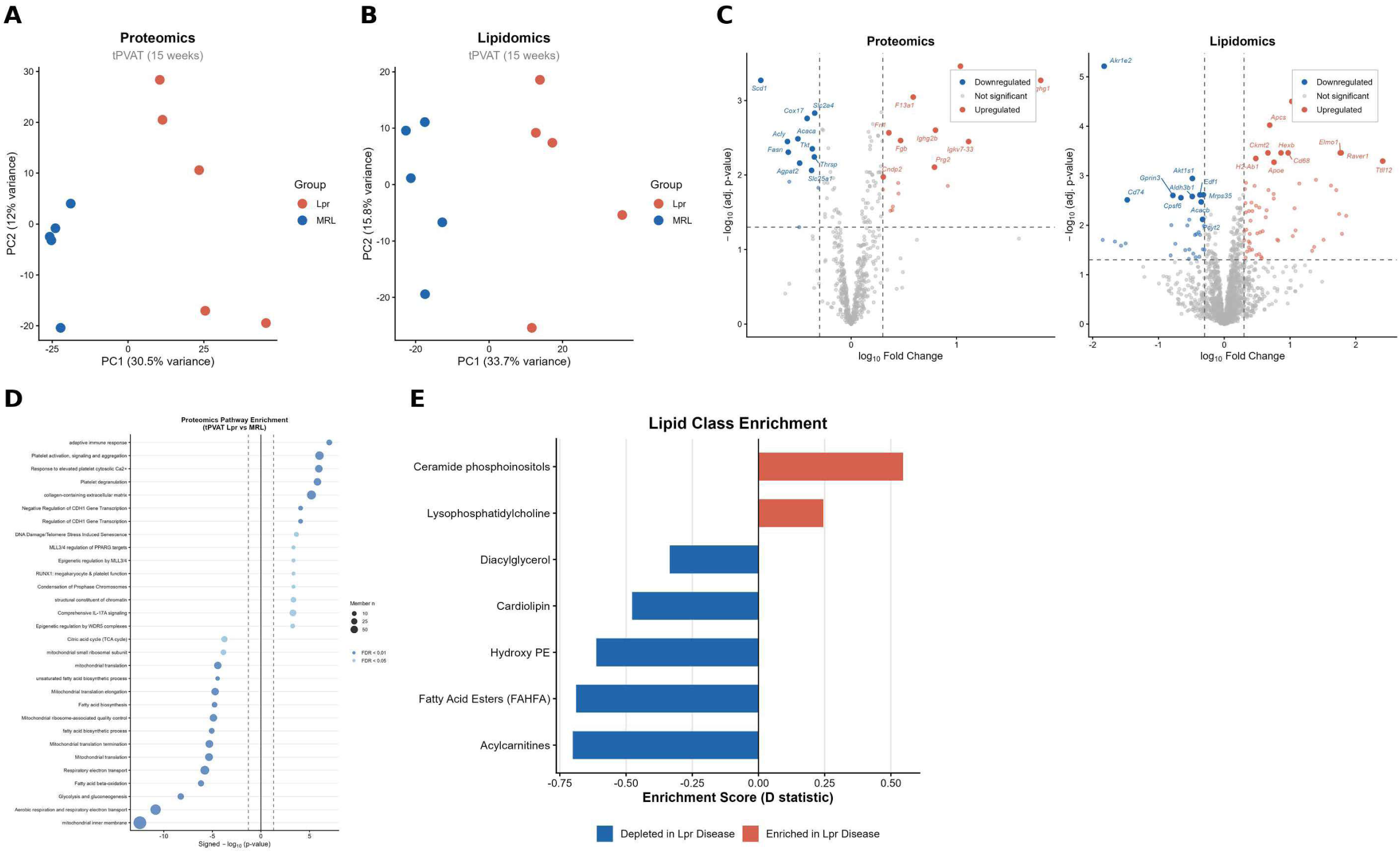
Multi-omics reveals coordinated metabolic suppression and lipidome remodeling in lupus tPVAT. (A,B) Principal component analysis (PCA) of (A) untargeted proteomics and (B) untargeted lipidomics from 15-week-old MRL (n = 5) and MRL/Lpr (n = 5) tPVAT, revealing distinct molecular signatures between groups. (C) Volcano plots highlighting differentially expressed proteins (left) and lipid species (right) in MRL/Lpr versus MRL tPVAT (adjusted p < 0.05; colored points meet significance thresholds), with selected features labeled. (D) Proteomics-based pathway enrichment analysis illustrating broad downregulation of mitochondrial bioenergetic pathways, including the TCA cycle, mitochondrial inner membrane components, and fatty acid β-oxidation, alongside enrichment of adaptive immune and platelet-associated pathways. Negative signed −log10(p value) indicates pathways decreased in MRL/Lpr, whereas positive values indicate pathways increased in MRL/Lpr. (E) Lipid class enrichment analysis showing depletion (negative D statistic) of acylcarnitines, FAHFAs, cardiolipin, hydroxy-PE species, and diacylglycerols, together with enrichment of ceramide phosphoinositols and lysophosphatidylcholines in MRL/Lpr tPVAT. Red bars indicate lipid classes enriched in MRL/Lpr, and blue bars indicate lipid classes depleted in MRL/Lpr.

### Multi-omics integration identifies convergent mitochondrial suppression and inflammatory niche formation in tPVAT from MRL/lpr mice

To determine whether transcriptomic, proteomic, and lipidomic changes converged on common biological programs, we performed integrated multi-omics analysis. A pathway concordance heatmap revealed coordinated suppression of mitochondrial and metabolic pathways, including mitochondrial translation, respiratory electron transport, the TCA cycle, fatty acid β-oxidation, fatty acid biosynthesis, and glycolysis, across RNA-seq, proteomics, and lipidomics (Figure 4A). In parallel, inflammatory pathways were enriched primarily across the transcriptomic and proteomic layers, including adaptive immune response, platelet activation, inflammatory response, and type I/II interferon-related signaling (Figure 4A). At the individual feature level, mitochondrial genes and proteins including *Echdc1, Slc25a1, Cox17, Sfxn1, Ldha, Cox6b1, Pkm,* and *Pdha1* were concordantly reduced across transcriptomic and proteomic datasets (Figure 4B). In contrast, several immune and antigen-presentation features were increased across the transcriptomic and proteomic layers. *Ighg1, Igkc, Ighg2b, H2-Aa, H2-Ab1,* and *Apoe* were elevated at both the mRNA and protein levels, whereas *Cd68* and *Clec10a* were increased at the protein level (Figure 4C). Global RNA-protein concordance analysis across 2,131 shared gene-protein pairs identified 128 concordantly upregulated and 27 concordantly downregulated pairs, with a smaller subset of discordant features (Figure 4D). Lipid-protein correlation analysis further showed that cardiolipin was strongly positively correlated with the mitochondrial/metabolic protein module and strongly inversely correlated with the immune/inflammatory protein module, whereas ceramides and lysophosphatidylcholines showed the opposite relationship (Figure 4E). Together, these data define a highly concordant cross-platform signature in tPVAT from MRL/lpr mice, characterized by mitochondrial and metabolic suppression, inflammatory activation, and coordinated lipid remodeling. (Figure 4F).

**Figure 4.**
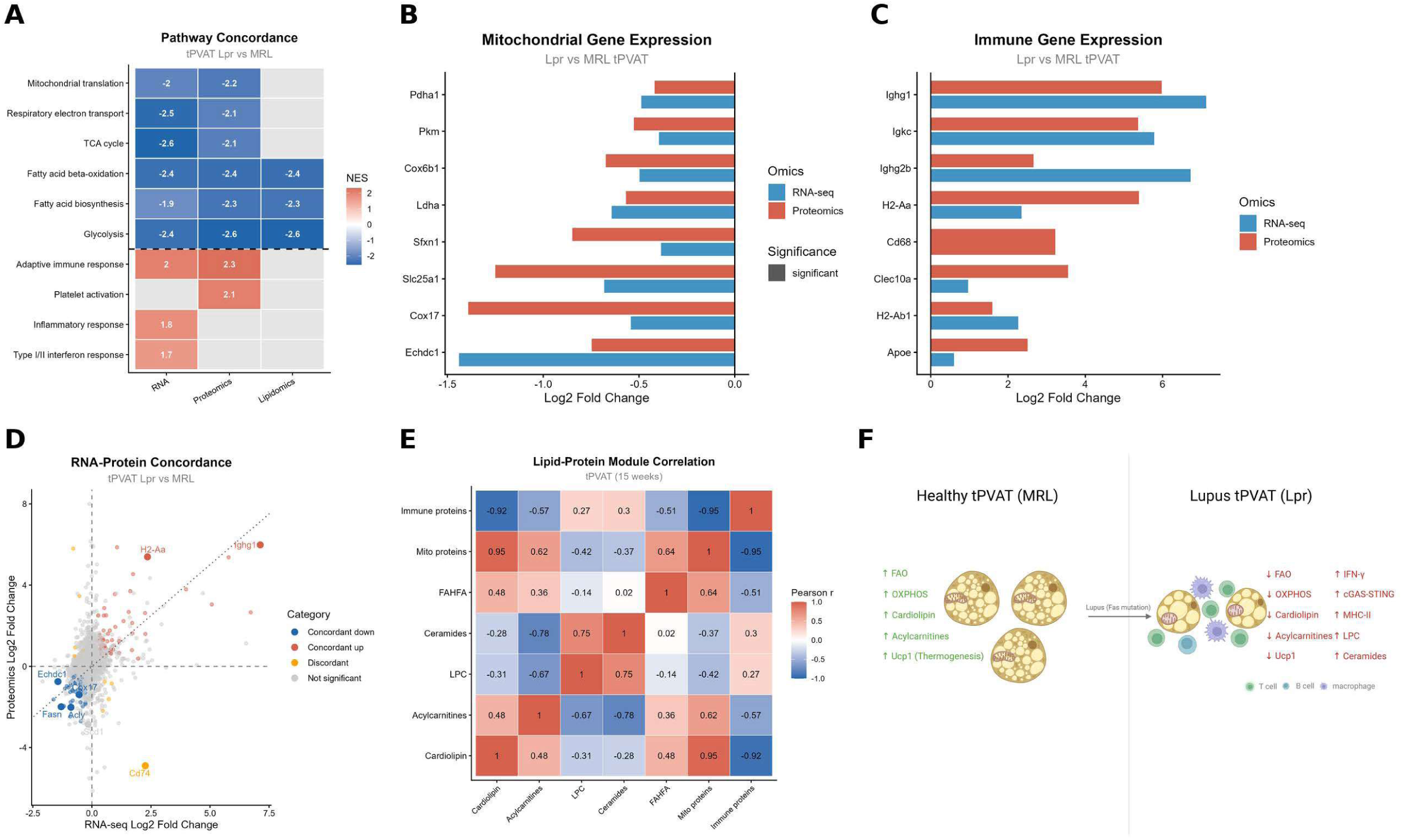
Integrated multi-omics reveals concordant collapse of mitochondrial bioenergetics and induction of an inflammatory niche. (A) Multi-omic concordance heatmap displaying normalized enrichment scores (NES) for selected pathways across transcriptomic (RNA-seq), proteomic, and lipidomic datasets, highlighting consistent suppression of metabolic pathways and activation of inflammatory signatures in MRL/Lpr versus MRL tPVAT. (B,C) Comparative log2 fold changes (MRL/Lpr vs MRL) demonstrating concordant (B) downregulation of mitochondrial respiratory chain and related metabolic genes/proteins and (C) increased abundance of selected immune and antigen-presentation markers across transcriptomic and proteomic layers. (D) Global RNA-protein concordance analysis in tPVAT; each point represents a gene-protein pair categorized as concordant upregulated, concordant downregulated, discordant, or not significant. (E) Pearson correlation matrix between lipid class summary scores and protein module scores, showing strong positive correlations between the mitochondrial/metabolic protein module and cardiolipin, FAHFAs, and acylcarnitines, and inverse correlations between these lipids and the immune/inflammatory protein module. Protein modules represent the mean log-transformed abundance of selected proteins, as defined in Methods. (F) Proposed working model: healthy MRL tPVAT maintains vascular protection through active fatty acid oxidation, oxidative phosphorylation, and Ucp1-driven thermogenesis with abundant cardiolipin and acylcarnitines, whereas lupus MRL/Lpr tPVAT exhibits immune accumulation, loss of cardiolipin-rich mitochondrial membranes, accumulation of ceramides and lysophosphatidylcholines, and failure of the thermogenic program.

### Lupus is associated with an acquired, depot-specific, and disease-stage-dependent defect in tPVAT ASPC adipogenesis

To determine whether the inflammatory and metabolic remodeling of PVAT from MRL/lpr mice is accompanied by altered progenitor cell function, we isolated ASPCs from 15-week-old MRL and MRL/lpr mice and assessed adipogenic differentiation under white and beige conditions by Oil Red O staining. ASPCs derived from tPVAT of MRL/lpr mice showed significantly reduced lipid accumulation under both white and beige differentiation conditions compared with MRL controls, and representative micrographs confirmed impaired lipid droplet formation under both conditions (Figure 5A, C). To determine whether this defect was present before overt disease progression, we repeated the assay using ASPCs isolated from 5-week-old mice. At this earlier time point, no significant genotype-dependent differences were observed under either white or beige differentiation conditions (Figure 5B, D), indicating that the adipogenic differentiation defect is acquired rather than developmentally fixed. To assess whether this impairment was depot-specific, we examined subcutaneous ASPCs from both 5-week-old and 15-week-old mice. In contrast to tPVAT-derived cells, subcutaneous ASPCs from MRL/lpr mice did not differ significantly from MRL controls under either white or beige differentiation conditions at either age (Figure S3). Together, these findings indicate that lupus selectively impairs the adipogenic potential of ASPCs from tPVAT at the active disease stage, consistent with an acquired, depot-specific progenitor defect.

**Figure 5.**
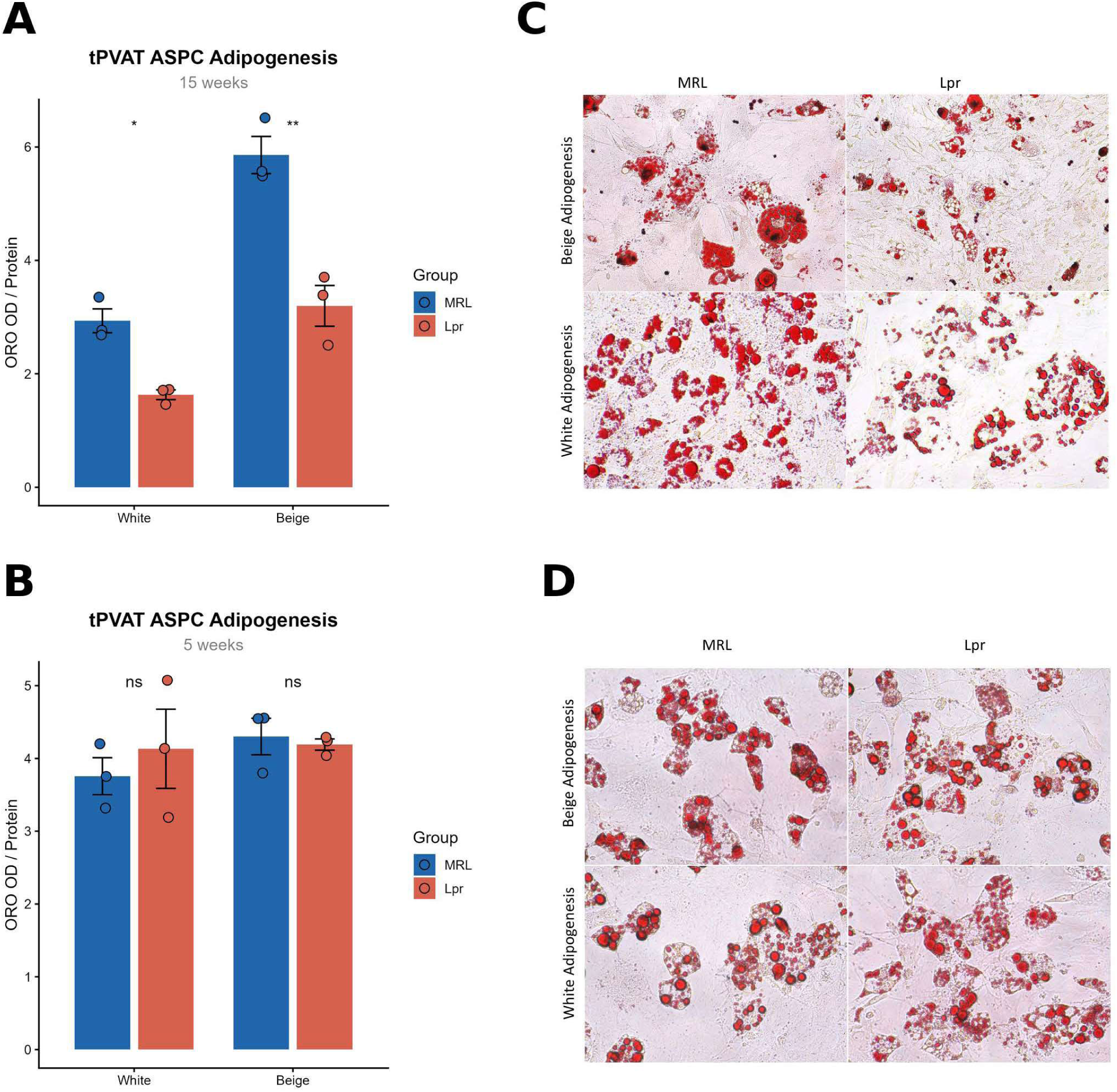
Lupus is associated with an acquired defect in tPVAT ASPC adipogenic differentiation. (A) Quantification of lipid accumulation (Oil Red O [ORO] absorbance normalized to total protein) in tPVAT adipose stem/progenitor cells (ASPCs) from 15-week-old MRL and MRL/Lpr mice cultured under white or beige adipogenic conditions. Data represent three independent biological experiments (n = 3), each using pooled tPVAT from three mice per genotype to ensure sufficient and representative progenitor populations. (B) ORO quantification in ASPCs isolated from 5-week-old mice, demonstrating preserved adipogenic potential in both genotypes at the pre-disease stage. (C, D) Representative ORO-stained micrographs of ASPC cultures from 15-week-old (C) and 5-week-old (D) mice under beige (top row) and white (bottom row) adipogenic differentiation conditions. Scale bar = 100 µm. Data are presented as mean ± SEM of the biological experiments; each biological replicate was assayed in technical triplicate. Statistical comparisons were performed by unpaired t-test separately for white and beige adipogenic conditions. *p<0.05, **p<0.01.

## Discussion

In this study, we provide the first integrated multi-omics characterization of tPVAT in a murine model of lupus, revealing coordinated cross-platform suppression of mitochondrial bioenergetic genes together with establishment of an inflammatory perivascular niche. Using bulk RNA-seq, untargeted proteomics, lipidomics, and high-dimensional spectral flow cytometry, we show that tPVAT from MRL/lpr mice undergoes broad transcriptomic, proteomic, and lipidomic remodeling that is highly concordant across platforms. We further demonstrate that lupus is associated with an acquired, depot-specific defect in ASPC adipogenic differentiation that emerges after disease development. Together, these findings substantially advance our understanding of the molecular basis of PVAT dysfunction in lupus and support a model in which immunometabolic reprogramming of the perivascular niche contributes to lupus-associated cardiovascular disease [22, 39].

### Mitochondrial dysfunction as a central feature of tPVAT in MRL/lpr mice

The most consistent finding across all three omics platforms was coordinated suppression of mitochondrial bioenergetic programs. At the transcriptomic level, fatty acid metabolism, oxidative phosphorylation, and adipogenesis were among the most strongly suppressed hallmark pathways. These findings were reinforced at the protein level, where mitochondrial inner membrane components, TCA cycle enzymes, fatty acid β-oxidation machinery, and the mitochondrial translation apparatus were all significantly reduced. At the lipid level, this mitochondrial dysfunction was accompanied by depletion of cardiolipin, a structural phospholipid required for inner mitochondrial membrane integrity, respiratory supercomplex stability, and efficient electron transport [40], as well as acylcarnitines, which reflect active fatty acid transport into mitochondria [41], and FAHFAs, which have been linked to anti-inflammatory and insulin-sensitizing effects [42]. The strong positive correlation between cardiolipin and the mitochondrial/metabolic protein module, together with their shared inverse relationship with the immune/inflammatory protein module, supports a close association between mitochondrial dysfunction and immune activation in lupus tPVAT.

This pattern is consistent with prior work demonstrating reduced *Ucp1* expression and adipocyte whitening in PVAT from mice with lupus [22], and extends those findings by showing a broader pattern of mitochondrial dysfunction. Notably, whereas *Ucp1* was significantly reduced, *Ucp2*, a stress-responsive uncoupling protein associated more with ROS buffering than adaptive thermogenesis, was selectively increased [43]. This pattern suggests reprogramming of tPVAT away from a specialized thermogenic state and toward a generalized stress-adaptive state that is unlikely to preserve normal vascular-supportive function. Similar shifts in mitochondrial uncoupling have been described in settings of oxidative stress and immune activation and may reflect a maladaptive response that further compromises mitochondrial homeostasis [43]. More broadly, these findings fit with emerging evidence that mitochondrial dysfunction in lupus can actively promote oxidative stress, immune activation, endothelial injury, and cardiovascular pathology, rather than simply arise as a downstream consequence [44]. The lipidomic changes observed here also fit within a broader lupus-associated metabolic context, as plasma lipidomics in patients with lupus identified reduced carnitines and widespread phospholipid remodeling associated with inflammation and cardiovascular risk [23].

### An IFN-dominant immune niche in tPVAT from MRL/lpr mice

At the transcriptomic level, the most strongly activated pathways in tPVAT from MRL/lpr mice were interferon-γ response, interferon-α response, IL6-JAK-STAT3 signaling, and TNFA signaling via NF-κB, collectively resembling the systemic type I/type II interferon signature that characterizes active SLE [45]. This IFN program was not only present at the RNA level but was also reflected at the protein level through concordant upregulation of MHC II-associated proteins (*H2-Aa, H2-Ab1*), myeloid markers (*Cd68, Clec10a, Apoe*), and immunoglobulin heavy and light chains (*Ighg1, Igkc, Ighg2b*), suggesting active antigen presentation and local humoral immune activity within the perivascular niche. A recent serum proteomics study similarly showed that lupus exhibits a particularly strong systemic proteomic signature relative to other autoimmune diseases, although our data extend this concept by localizing immune activation specifically to tPVAT [46].

Flow cytometry defined the cellular basis of this signature. tPVAT from MRL/lpr mice harbored marked expansion of CD45+ leukocytes driven predominantly by T cells. Both CD4+ and CD8+ T cells were significantly elevated, with preferential accumulation of CD44−CD62L− cells in both compartments, indicating expansion of activated, non-naive T-cell states within the perivascular niche. Although Tregs increased in absolute number, the Treg/CD4+ ratio was significantly reduced, indicating incomplete immunoregulatory compensation. This disruption of the Treg/effector balance within the perivascular niche is particularly relevant because Tregs are known to suppress endothelial activation and vascular inflammation [47]. Prior work showed that PVAT from MRL/lpr mice promotes endothelial dysfunction in ex vivo vascular assays, and the immunological data presented here suggest that local loss of immunoregulatory balance may represent one cellular mechanism contributing to that dysfunction [22].

The lipidomic data add an additional mechanistic layer. Accumulation of ceramide phosphoinositols and LPCs in tPVAT from MRL/lpr mice points toward a pro-inflammatory lipid microenvironment. LPCs are well-established mediators of endothelial activation, monocyte recruitment, and NF-κB signaling [48, 49]. Ceramides, in turn, have been linked to insulin resistance, apoptosis, oxidative stress, mitochondrial dysfunction, and inflammatory signaling [50, 51]. Both lipid classes have also been implicated in endoplasmic reticulum stress and mitochondrial reactive oxygen species production in metabolic tissues [48, 50, 52]. The inverse correlation between these pro-inflammatory lipid classes and mitochondrial protein modules suggests that lipid remodeling and mitochondrial dysfunction are closely coupled in tPVAT from MRL/lpr mice and may reinforce one another within the perivascular niche. The broader lupus lipidomics literature also supports an association between lipid remodeling, inflammation, and cardiovascular risk, although the direction of several individual lipid classes differs between plasma in humans and our tPVAT dataset in mice [23].

### An acquired, depot-specific progenitor defect as a functional consequence of lupus

A key novel finding of this study is that lupus is associated with a functional defect in resident perivascular ASPCs. ASPCs isolated from tPVAT of15-week-old MRL/lpr mice showed significantly impaired lipid accumulation under both white and beige differentiation conditions compared with MRL controls, whereas no significant genotype-dependent differences were observed at 5 weeks of age. The specificity of this defect, absent at the pre-disease time point and confined to the tPVAT depot, with subcutaneous ASPCs remaining fully competent at both ages, indicates that this is not a constitutive progenitor abnormality but rather an acquired, disease-stage- and niche-dependent phenotype. Taken together, the convergence of bulk multi-omics and functional progenitor data supports the idea that the inflamed perivascular niche, rather than intrinsic genetic differences, contributes to impaired adipogenesis and loss of thermogenic phenotype in lupus tPVAT.

This observation has important implications for vascular biology. Under physiological conditions, PVAT is maintained in part through local progenitor renewal, which supports the pool of thermogenically competent adipocytes linked to vascular homeostasis [53, 54]. Healthy ASPCs are capable of self-renewal and adipogenic differentiation to generate new, mature adipocytes required for maintenance of PVAT tissue function, and thoracic PVAT is recognized as containing UCP1-positive brown/beige adipocytes that contribute to vascular homeostasis. Suppression of such functions in lupus would thus be expected to accelerate PVAT whitening and functional deterioration, thereby compounding its pro-inflammatory and vasoconstrictive properties [22, 55, 56]. Consistent with this idea, brown adipose involution has been linked to progressive restriction in progenitor competence, and dysfunctional or inflamed PVAT has been associated with vascular dysfunction and loss of vaso-protective properties. Our data therefore support a feed-forward model in which immune infiltration, mitochondrial dysfunction, and lipid remodeling create a microenvironment that suppresses ASPC differentiation, limits thermogenic replenishment, and perpetuates PVAT dysfunction, a process that may become increasingly difficult to reverse at later disease stages.

### Limitations and future directions

Several limitations should be acknowledged. First, this study focused on a single disease time point at 15 weeks, and the temporal sequence of transcriptomic, proteomic, lipidomic, and cellular changes, as well as their causal relationships, remains to be defined through longitudinal multi-omics profiling. Second, although we demonstrated an acquired defect in ASPC adipogenic differentiation in vitro, the specific inflammatory and metabolic signals within the lupus perivascular niche, such as interferons, ceramides, LPCs, or direct T cell-derived cues, that mediate progenitor suppression have not yet been identified and represent an important focus for future mechanistic studies. Third, although MRL/lpr mice are a well-established model of lupus, the Fas mutation that drives lymphoproliferation may not fully recapitulate all aspects of human lupus; accordingly, the relevance of these findings to human thoracic PVAT pathology will need to be tested in future studies using patient imaging, blood-based profiling, and tissue samples. Finally, the functional consequences of these multi-omic changes for endothelial function, vascular stiffness, and atherosclerotic lesion formation in vivo remain to be determined directly through pathway-targeted intervention studies. Future work should test whether strategies aimed at restoring mitochondrial function, dampening interferon signaling, or modifying pathogenic lipid remodeling can restore the tPVAT progenitor niche and preserve the beige, vasculoprotective PVAT phenotype in lupus-prone mice.

## Conclusions

This study provides the first integrated multi-omics portrait of lupus tPVAT dysfunction, demonstrating a concordant cross-platform diminution of mitochondrial bioenergetics, a shift toward pro-inflammatory and lipotoxic lipid remodeling, marked T cell-dominant immune infiltration with loss of immunoregulatory balance, and an acquired, depot-specific defect in ASPC adipogenic capacity. Together, these findings establish a molecular framework for understanding how the lupus perivascular niche loses its vasculoprotective identity and acquires pathogenic features that may contribute to the accelerated cardiovascular disease observed in lupus. Targeting the immunometabolic dysfunction of PVAT, through restoration of mitochondrial bioenergetics, suppression of the interferon-dominant immune niche, or rescue of progenitor differentiation capacity may represent a promising therapeutic strategy for lupus-associated cardiovascular disease.

## Declarations

### Ethics approval and consent to participate

All animal care and experimental procedures were conducted in accordance with the National Institutes of Health Guide for the Care and Use of Laboratory Animals and relevant institutional guidelines. All protocols were approved by the Institutional Animal Care and Use Committee of Augusta University.

### Consent for publication

Not applicable.

### Availability of data and materials

The datasets generated and/or analyzed during the current study are available from the corresponding author on reasonable request. Relevant source data are included in the article and its supplementary information files.

### Authors’ contributions

HS and NLW conceived and designed the study. HS, LLL, DK, YZ, BG, XX, TZ, QH, QL and MO performed experiments. HS, LLL, XX and SW analyzed and interpreted the data. SW and HS prepared the figures. HS, SW and NLW drafted the manuscript. HS, NLW, HPW, VP, DG, RL, HWK, DS, DF, JMK, LC, KL and BHA critically revised the manuscript for important intellectual content. All authors read and approved the final manuscript.

## Acknowledgements

The authors thank Georgia Cancer Center Flow and Mass Cytometry Core Facility (RRID: SCR_025747) for support with flow cytometry. The authors also thank Dalton Bioanalytics for proteomics and lipidomics analysis and Azenta Life Sciences for RNA sequencing services.

---

**Supplemental Figure S1.**
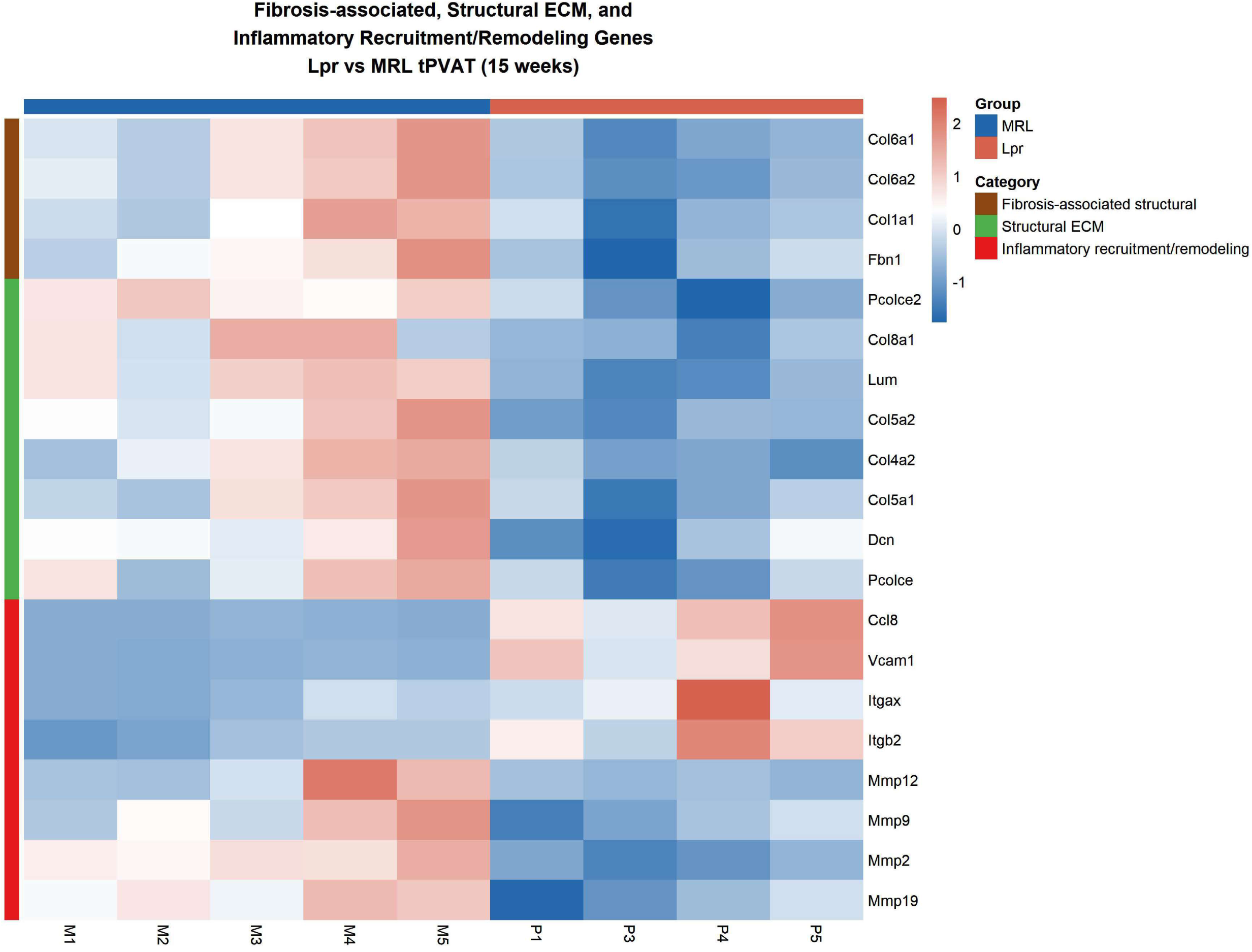
Fibrosis-associated structural, ECM, and inflammatory recruitment/remodeling gene expression in lupus tPVAT. Heatmap of selected fibrosis-associated structural genes, structural ECM genes, and inflammatory recruitment/remodeling genes in tPVAT from 15-week-old MRL (n = 5) and MRL/Lpr (n = 4) mice. Expression values are shown as row-scaled normalized bulk RNA-seq expression across samples. MRL/Lpr tPVAT demonstrated reduced expression of multiple fibrosis-associated structural and ECM genes, together with increased expression of inflammatory recruitment genes, including *Ccl8, Vcam1, Itgax,* and *Itgb2*. Matrix metalloproteinase genes (*Mmp12, Mmp9, Mmp2,* and *Mmp19*) were reduced in MRL/Lpr, consistent with loss of homeostatic stromal matrix programs rather than uniform activation of a canonical fibrotic transcriptional response. Color scale represents row-scaled normalized counts (z-scores).

**Supplemental Figure S2.**
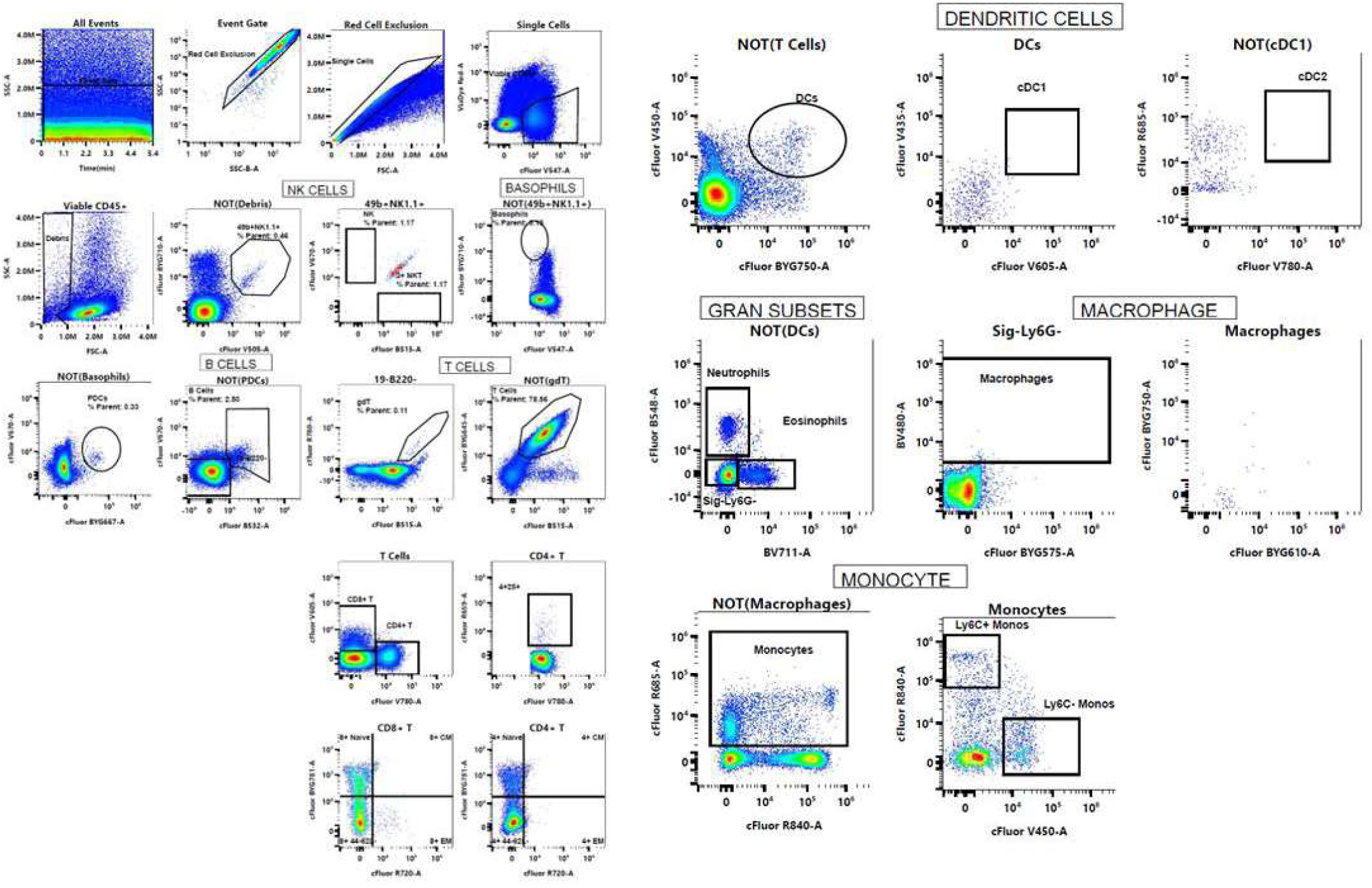
Spectral flow cytometry gating strategy for immune cell analysis in tPVAT. Representative gating strategy used to identify viable CD45+ leukocytes and downstream immune populations from tPVAT. Sequential gates included event quality control, red cell exclusion, singlet discrimination, and live CD45+ cells, followed by identification of NK cells, basophils, B cells, T cells, dendritic cells, granulocyte subsets, macrophages, and monocytes. T cells were further subdivided into CD4+, CD8+, and γδ T cells, and CD4+ and CD8+ T cells were further classified into naïve (CD44−CD62L+), central memory (CM; CD44+CD62L+), effector memory (EM; CD44+CD62L−), and CD44−CD62L− atypical/pre-effector-like subsets. Dendritic cells were further resolved into cDC1 and cDC2 populations, and monocytes into Ly6C+ and Ly6C− subsets. Representative plots are shown from one sample; the same gating strategy was applied across all animals.

**Supplemental Figure S3.**
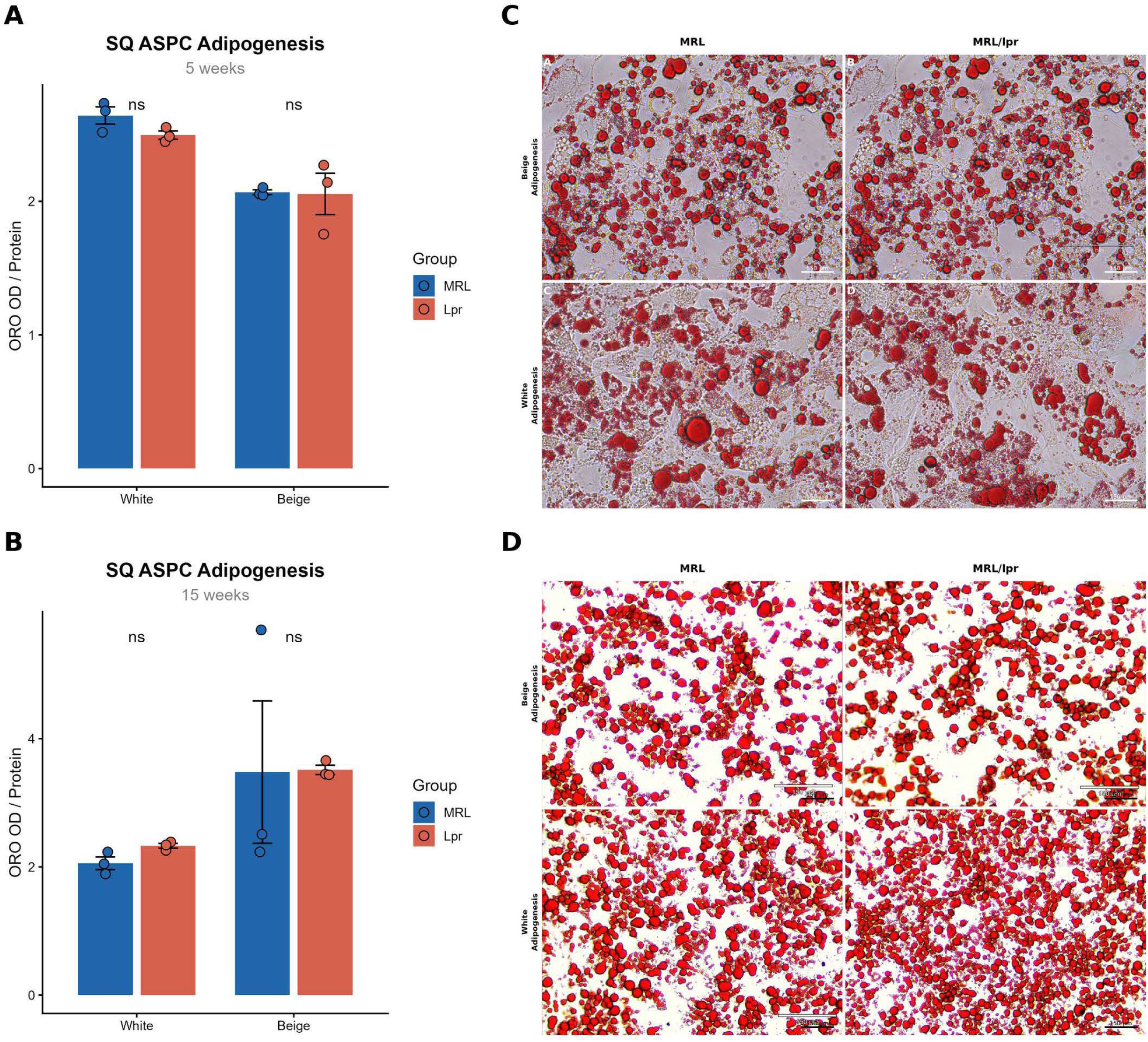
Preservation of subcutaneous ASPC adipogenic potential in lupus mice. (A,B) Quantification of lipid accumulation (Oil Red O [ORO] absorbance normalized to total protein) in subcutaneous (SQ) adipose stem/progenitor cells (ASPCs) isolated from MRL and MRL/Lpr mice at (A) 5 weeks and (B) 15 weeks of age, differentiated under white or beige adipogenic conditions. In contrast to the tPVAT depot, SQ ASPCs did not differ significantly between genotypes under either differentiation condition at either age. (C,D) Representative ORO-stained micrographs of SQ ASPC cultures from (C) 5-week-old and (D) 15-week-old cohorts following beige (top row) and white (bottom row) adipogenic differentiation. Scale bar = 150 µm. Data are presented as mean ± SEM of three independent biological experiments, each using pooled SQ ASPCs from three mice per genotype; each biological replicate was assayed in technical triplicate. Statistical comparisons were performed by unpaired t-test separately for white and beige adipogenic conditions. ns = not significant.

## References

1. Tsokos, G.C., Systemic lupus erythematosus. N Engl J Med, 2011. 365(22): p. 2110–21.

2. Oliveira, C.B. and M.J. Kaplan, Cardiovascular disease risk and pathogenesis in systemic lupus erythematosus. Semin Immunopathol, 2022. 44(3): p. 309–324.

3. Manzi, S., et al., Age-specific incidence rates of myocardial infarction and angina in women with systemic lupus erythematosus: comparison with the Framingham Study. Am J Epidemiol, 1997. 145(5): p. 408–15.

4. Joyce, D.P., et al., Prevalence of cardiovascular events in a population-based registry of patients with systemic lupus erythematosus. Arthritis Res Ther, 2024. 26(1): p. 160.

5. Moghaddam, B., et al., All-cause and cause-specific mortality in systemic lupus erythematosus: a population-based study. Rheumatology (Oxford), 2021. 61(1): p. 367–376.

6. Ajeganova, S., I. Hafstrom, and J. Frostegard, Patients with SLE have higher risk of cardiovascular events and mortality in comparison with controls with the same levels of traditional risk factors and intima-media measures, which is related to accumulated disease damage and antiphospholipid syndrome: a case-control study over 10 years. Lupus Sci Med, 2021. 8(1).

7. McMahon, M., B.H. Hahn, and B.J. Skaggs, Systemic lupus erythematosus and cardiovascular disease: prediction and potential for therapeutic intervention. Expert Rev Clin Immunol, 2011. 7(2): p. 227–41.

8. McMahon, M. and B.J. Skaggs, Rethinking Cardiovascular Screening in Systemic Lupus Erythematosus. J Rheumatol, 2026.

9. Esdaile, J.M., et al., Traditional Framingham risk factors fail to fully account for accelerated atherosclerosis in systemic lupus erythematosus. Arthritis Rheum, 2001. 44(10): p. 2331–7.

10. Toloza, S.M., et al., Systemic lupus erythematosus in a multiethnic US cohort (LUMINA): XXII. Predictors of time to the occurrence of initial damage. Arthritis Rheum, 2004. 50(10): p. 3177–86.

11. Woodridge, L., et al., Subclinical Atherosclerosis Risk Can Be Predicted in Female Patients With Systemic Lupus Erythematosus Using Metabolomic Signatures: An Observational Study. J Am Heart Assoc, 2025. 14(8): p. e036507.

12. Fernandez-Alfonso, M.S., et al., Role of PVAT in coronary atherosclerosis and vein graft patency: friend or foe? Br J Pharmacol, 2017. 174(20): p. 3561–3572.

13. Kim, H.W., et al., Perivascular Adipose Tissue and Vascular Perturbation/Atherosclerosis. Arterioscler Thromb Vasc Biol, 2020. 40(11): p. 2569–2576.

14. Mancio, J., E.K. Oikonomou, and C. Antoniades, Perivascular adipose tissue and coronary atherosclerosis. Heart, 2018. 104(20): p. 1654–1662.

15. Horimatsu, T., et al., Remote Effects of Transplanted Perivascular Adipose Tissue on Endothelial Function and Atherosclerosis. Cardiovasc Drugs Ther, 2018. 32(5): p. 503–510.

16. Gu, W., et al., Adventitial Cell Atlas of wt (Wild Type) and ApoE (Apolipoprotein E)-Deficient Mice Defined by Single-Cell RNA Sequencing. Arterioscler Thromb Vasc Biol, 2019. 39(6): p. 1055–1071.

17. Koenen, M., et al., Ablation of Prdm16 and beige fat causes vascular remodeling and elevated blood pressure. bioRxiv, 2025.

18. Pan, X., et al., SM22alpha-Lineage Perivascular Stromal Cells Contribute to Abdominal Aortic Aneurysm. Circ Res, 2025. 137(1): p. 4–22.

19. Shields, K.J., et al., Perivascular adipose tissue of the descending thoracic aorta is associated with systemic lupus erythematosus and vascular calcification in women. Atherosclerosis, 2013. 231(1): p. 129–35.

20. Shields, K.J., et al., Association of aortic perivascular adipose tissue density with aortic calcification in women with systemic lupus erythematosus. Atherosclerosis, 2017. 262: p. 55–61.

21. Roldan, L.P., et al., Aortic adventitial thickness as a marker of aortic atherosclerosis, vascular stiffness, and vessel remodeling in systemic lupus erythematosus. Clin Rheumatol, 2021. 40(5): p. 1843–1852.

22. Shi, H., et al., Perivascular adipose tissue promotes vascular dysfunction in murine lupus. Front Immunol, 2023. 14: p. 1095034.

23. Duarte-Delgado, N.P., et al., Metabolic Alterations in Colombian Women with Rheumatoid Arthritis and Systemic Lupus Erythematosus Reveal Potential Lipid Biomarkers Associated with Inflammation and Cardiovascular Risk. Int J Mol Sci, 2025. 26(10).

24. Crow, M.K., Type I interferon in the pathogenesis of lupus. J Immunol, 2014. 192(12): p. 5459–68.

25. Pilkington, A.C., H.A. Paz, and U.D. Wankhade, Beige Adipose Tissue Identification and Marker Specificity-Overview. Front Endocrinol (Lausanne), 2021. 12: p. 599134.

26. Hass, D.T. and C.J. Barnstable, Uncoupling proteins in the mitochondrial defense against oxidative stress. Prog Retin Eye Res, 2021. 83: p. 100941.

27. Nesci, S. and S. Rubattu, UCP2, a Member of the Mitochondrial Uncoupling Proteins: An Overview from Physiological to Pathological Roles. Biomedicines, 2024. 12(6).

28. Cox, A.R., et al., STAT1 Dissociates Adipose Tissue Inflammation From Insulin Sensitivity in Obesity. Diabetes, 2020. 69(12): p. 2630–2641.

29. Deng, Y., et al., Expression characteristics of interferon-stimulated genes and possible regulatory mechanisms in lupus patients using transcriptomics analyses. EBioMedicine, 2021. 70: p. 103477.

30. Kim, B.H., et al., IFN-inducible GTPases in host cell defense. Cell Host Microbe, 2012. 12(4): p. 432–44.

31. Bradley, D., et al., Interferon gamma mediates the reduction of adipose tissue regulatory T cells in human obesity. Nat Commun, 2022. 13(1): p. 5606.

32. Sun, K., X. Li, and P.E. Scherer, Extracellular Matrix (ECM) and Fibrosis in Adipose Tissue: Overview and Perspectives. Compr Physiol, 2023. 13(1): p. 4387–4407.

33. Chung, K.J., et al., A self-sustained loop of inflammation-driven inhibition of beige adipogenesis in obesity. Nat Immunol, 2017. 18(6): p. 654–664.

34. Wu, H., et al., CD11c expression in adipose tissue and blood and its role in diet-induced obesity. Arterioscler Thromb Vasc Biol, 2010. 30(2): p. 186–92.

35. Wu, Y., et al., FAP expression in adipose tissue macrophages promotes obesity and metabolic inflammation. Proc Natl Acad Sci U S A, 2023. 120(51): p. e2303075120.

36. Ruiz-Ojeda, F.J., et al., Extracellular Matrix Remodeling of Adipose Tissue in Obesity and Metabolic Diseases. Int J Mol Sci, 2019. 20(19).

37. Li, W., et al., The Regulatory T Cell in Active Systemic Lupus Erythematosus Patients: A Systemic Review and Meta-Analysis. Front Immunol, 2019. 10: p. 159.

38. Parietti, V., et al., Function of CD4+,CD25+ Treg cells in MRL/lpr mice is compromised by intrinsic defects in antigen-presenting cells and effector T cells. Arthritis Rheum, 2008. 58(6): p. 1751–61.

39. Adachi, Y., K. Ueda, and E. Takimoto, Perivascular adipose tissue in vascular pathologies-a novel therapeutic target for atherosclerotic disease? Front Cardiovasc Med, 2023. 10: p. 1151717.

40. Paradies, G., et al., Role of Cardiolipin in Mitochondrial Function and Dynamics in Health and Disease: Molecular and Pharmacological Aspects. Cells, 2019. 8(7).

41. Longo, N., M. Frigeni, and M. Pasquali, Carnitine transport and fatty acid oxidation. Biochim Biophys Acta, 2016. 1863(10): p. 2422–35.

42. Yore, M.M., et al., Discovery of a class of endogenous mammalian lipids with anti-diabetic and anti-inflammatory effects. Cell, 2014. 159(2): p. 318–32.

43. Brand, M.D. and T.C. Esteves, Physiological functions of the mitochondrial uncoupling proteins UCP2 and UCP3. Cell Metab, 2005. 2(2): p. 85–93.

44. Wang, H., et al., Mitochondria dysfunction: A trigger for cardiovascular diseases in systemic lupus erythematosus. Int Immunopharmacol, 2025. 144: p. 113722.

45. Ronnblom, L., Systemic lupus erythematosus: from an adverse event of interferon administration to a disease with new treatment options. Ups J Med Sci, 2025. 130.

46. Hocaoglu, M., J. Das, and A.H. Sawalha, Identifying systemic lupus erythematosus from serum proteomic profiles using machine learning and genetic risk stratification. Arthritis Rheumatol, 2026.

47. Zammit, N.W., et al., Regulatory T cells in brown adipose tissue safeguard thermogenesis by restraining interferon-gamma-producing lymphocytes. Sci Immunol, 2025. 10(109): p. eads0478.

48. Liu, P., et al., The mechanisms of lysophosphatidylcholine in the development of diseases. Life Sci, 2020. 247: p. 117443.

49. Bi, X., et al., Activation of liver X receptor attenuates lysophosphatidylcholine-induced IL-8 expression in endothelial cells via the NF-kappaB pathway and SUMOylation. J Cell Mol Med, 2016. 20(12): p. 2249–2258.

50. Fucho, R., et al., Ceramides and mitochondrial fatty acid oxidation in obesity. FASEB J, 2017. 31(4): p. 1263–1272.

51. Pan, Y., et al., A review of the mechanisms of abnormal ceramide metabolism in type 2 diabetes mellitus, Alzheimer’s disease, and their co-morbidities. Front Pharmacol, 2024. 15: p. 1348410.

52. de la Monte, S.M., et al., Dysfunctional pro-ceramide, ER stress, and insulin/IGF signaling networks with progression of Alzheimer’s disease. J Alzheimers Dis, 2012. 30 Suppl 2(02): p. S217–29.

53. Lin, T., et al., Mechanisms and metabolic consequences of adipocyte progenitor replicative senescence. Immunometabolism (Cobham), 2024. 6(3): p. e00046.

54. Brown, N.K., et al., Perivascular adipose tissue in vascular function and disease: a review of current research and animal models. Arterioscler Thromb Vasc Biol, 2014. 34(8): p. 1621–30.

55. Huang, Z., et al., Brown adipose tissue involution associated with progressive restriction in progenitor competence. Cell Rep, 2022. 39(2): p. 110575.

56. Nosalski, R. and T.J. Guzik, Perivascular adipose tissue inflammation in vascular disease. Br J Pharmacol, 2017. 174(20): p. 3496–3513.

